# 3D Genome Contributes to Protein-Protein Interactome

**DOI:** 10.1101/2020.11.30.404517

**Authors:** Yi Shi, Song Cao, Mingxuan Zhang, Xianbin Su, Zehua Guo, Jiayi Wang, Tingting Liu, Hui Shi, Guang He

## Abstract

Numerous computational methods have been proposed to predict protein-protein interactions, none of which however, considers the original DNA loci of the interacting proteins in the perspective of 3D genome. Here we retrospect the DNA origins of the interacting proteins in the context of 3D genome and discovered that 1) if a gene pair is more proximate in 3D genome, their corresponding proteins are more likely to interact. 2) signal peptide involvement of PPI affects the corresponding gene-gene proximity in 3D genome space. 3) by incorporating 3D genome information, existing PPI prediction methods can be further improved in terms of accuracy. Combining our previous discoveries, we conjecture the existence of 3D genome driven cellular compartmentalization, meaning the co-localization of DNA elements lead to increased probability of the co-localization of RNA elements and protein elements.

## Introduction

In almost all cellular processes, including DNA transcription and replication, signaling cascades, metabolic cycles and many additional processes, proteins undertake their cellular functions by coordinating with other proteins^1^. It is therefore important to know the specific nature of these protein-protein interactions (PPIs). A human cell at any time, contains over 100,000 binary interactions between proteins^2^, a small fraction of these protein-protein interactions however, are experimentally identified^3^, lagging behind the generation of sequencing information which grow exponentially. Biological-wise, this is due to the dynamic nature of these interactions, that many of them are transient, and others occur only in certain cellular contexts or at particular times in development^3^. Technology-wise, this is due to the low throughput and inherent imperfection of the empirical PPI identification experiments; for example, yeast two-hybrid (Y2H) system^4^ and co-immunoprecipitation (coIP) coupled with mass spectrometry^5^ are two widely adopted methods, both prone to false discoveries because procedures from the reagent choosing to the cell type used and experimental conditions can all influence the final outcome^6^.

To bridge the gap between ensemble in situ PPI and the identified ones, accurate and efficient computational methods are required, as the prediction results can either be directly used or boost the labor-intensive empirical methods. In the past two decades, numerous computational protein interaction discovery approaches have been developed. A PPI prediction method is usually determined by two factors: the first factor is the encoding scheme, i.e., what information is adopted and how they are encoded for the target protein or protein pair; the other factor is the mathematical learning model being employed. By combining these two factors, computational PPI prediction approaches can be further categorized into four classes: network topology based, genomic context and structural information based, text mining based, and machine learning based which utilize heterogeneous genomic or proteomic features. Many studies have demonstrated that utilizing these PPI prediction tools is important for new research in protein-protein interaction analysis to be conducted^1,7–11^.

It has been reported that genes that are proximate to each other in terms of linear genomic distance, could lead to their protein counterparts interacting to each other^12^. This occurs to us that genes that are proximate in 3D genomic space may also obey such rule, and chromatin conformation capturing technologies such as Hi-C^13,14^ and ChIA-PET^15^ developed in recent years provide an excellent opportunity to systematically investigate this conjecture. To the best of our knowledge, there is no existing PPI prediction method that considers the genomic 3D distance of the corresponding gene pairs so far. Therefore, if the gene-gene 3D distances are indeed correlated to the protein-protein interaction, it would contribute to the PPI prediction without doubt.

In this work, we retrospect the DNA origins of the interacting proteins in the context of 3D genome and discovered that 1) if a gene pair is more proximate in 3D genome, their corresponding proteins are more likely to interact. 2) signal peptide involvement of PPI affects the corresponding gene-gene proximity in 3D genome space. 3) by incorporating 3D genome information, existing PPI prediction methods can be further improved in terms of accuracy. Furthermore, by combining our previous discoveries – that somatic co-mutation DNA loci tend to form Somatic Co-mutation Hotspots (SCHs) in 3D genome space^16^, which was recently supported by Akdemir *et al*.^17^, and that 3D genome contribute to immunogenic neoantigen distribution^18^ – we conjecture the existence of 3D genome driven cellular compartmentalization; with this compartmentalization, the co-localization of DNA elements lead to increased probability of the co-localization of their downstream elements including RNAs, proteins, and even metabolic molecules, as Figure 1 illustrates.

**Figure 1.**
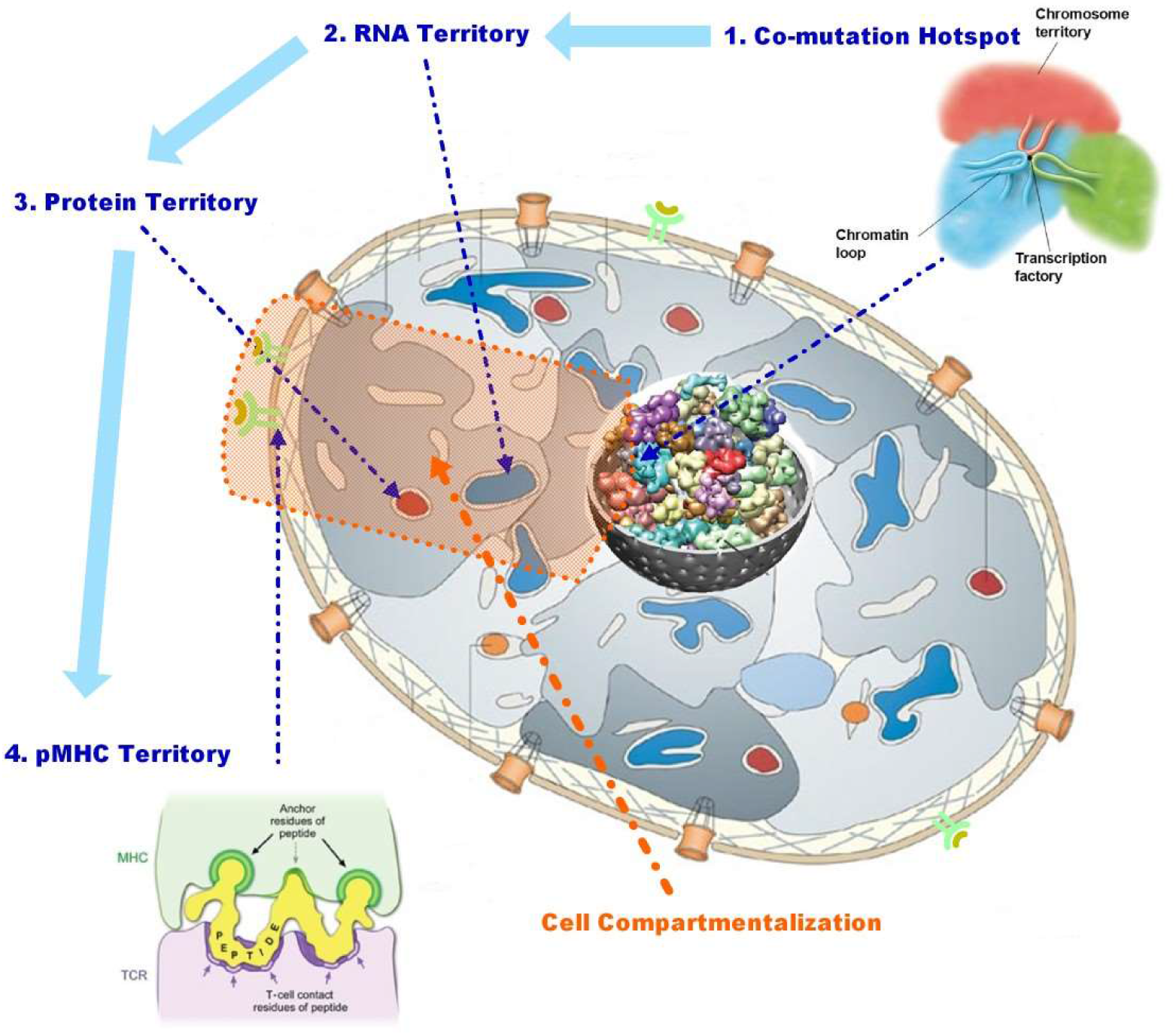
Conceptual illustration of 3D genome driven cell compartmentalization. Colocalization of DNA lead to co-expression of RNA, which further lead to protein-protein interaction and concentration of depredated molecular.

## Results

### Protein-protein interaction and 3D genome

To investigate whether the interacting proteins’ corresponding genes are more proximate to each other in chromatin 3D space, we conducted intra-chromosomal (per each individual chromosome) and inter-chromosomal (whole genome) analyses. For the intra-chromosomal analyses, we compared PPIs’ corresponding gene-gene spatial contact frequencies (inverse to 3D distance) on Hi-C heatmaps with the overall background and gene-level background contact frequencies of DNA loci of the same linear distance. As Figure 2 demonstrates, the PPIs’ gene counterparts are significantly more proximate to each other comparing to both background values.

**Figure 2.**
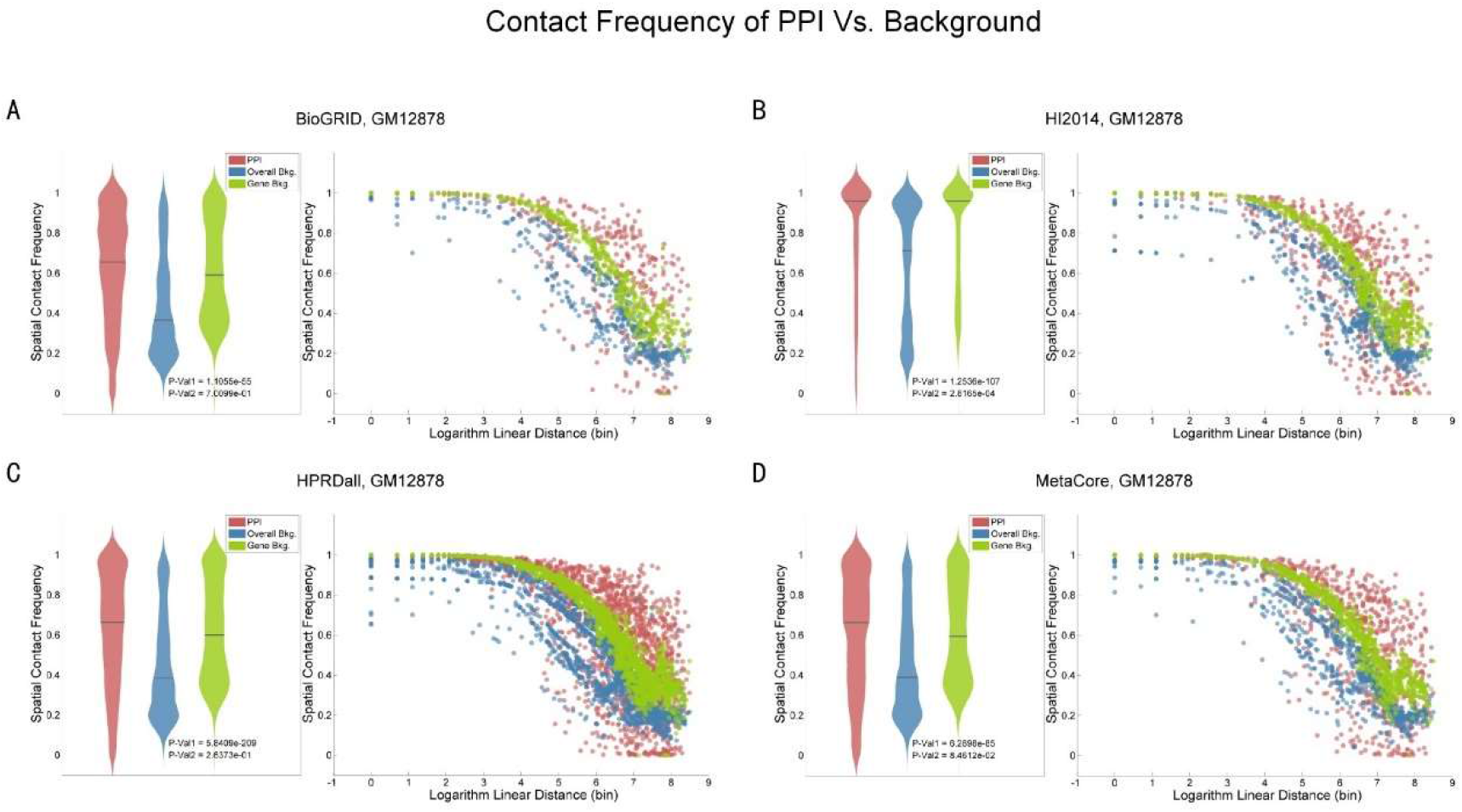
Comparison of chromatin contact frequency (inverse of 3D distance) distributions of PPI and background in intra-chromosomal level. Left: PPIs’ corresponding gene pair 3D distance distribution (red), background 3D distance distribution with same linear distance (blue), gene-level background 3D distance distribution with same linear distance (green). Right: detailed scatter points of 3D distances (y-axis) along with linear distances (x-axis). A,B,C,D correspond to BioGRID, HI2014, HPRDall, and MetaCore PPI databases analyzed with GM12878 Hi-C cell line.

For the inter-chromosomal analyses, we first generated non-PPI pairs for each PPI dataset so that each non-PPI protein pair is never witnessed be interacting by any previous empirical experiment. We then compare the corresponding gene-gene contact frequencies and the neighboring regions of both PPI and non-PPI. As Figure 3 and Figure 4 demonstrate, the PPIs’ corresponding gene-gene contact frequencies including their neighboring regions are significantly more proximate to each other compared with the non-PPIs’ gene-gene pairs.

**Figure 3.**
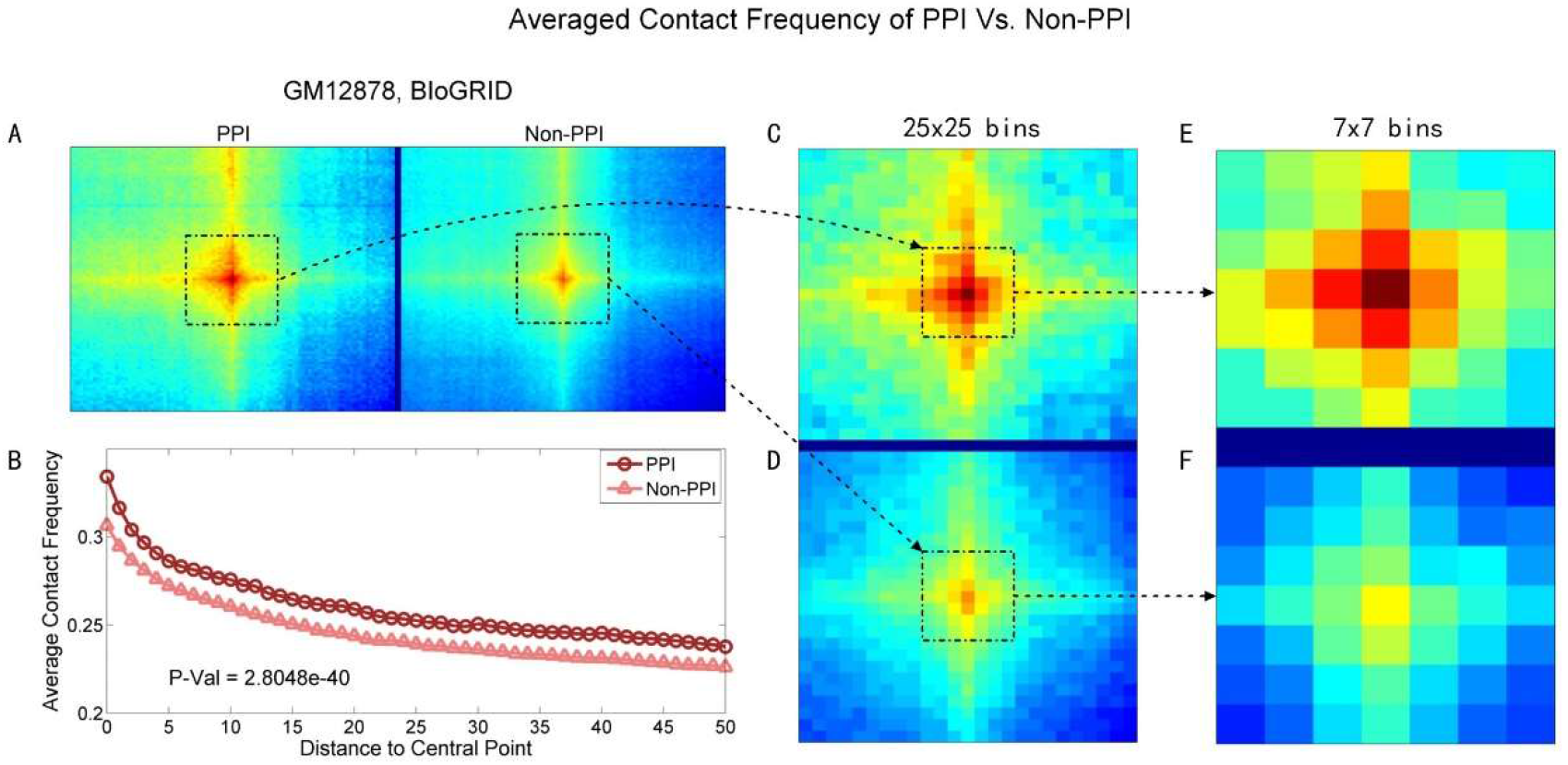
A: averaged Hi-C heatmap (101×101 bins) centered at PPIs’ and Non-PPIs’ corresponding gene-gene loci. B: PPIs and Non-PPIs’ corresponding averaged contact frequencies and their decay along with increased distance to the central point. C, D: zoomed in heatmap with 25×25 bins. E, F: zoomed in heatmap with 7×7 bins.

**Figure 4.**
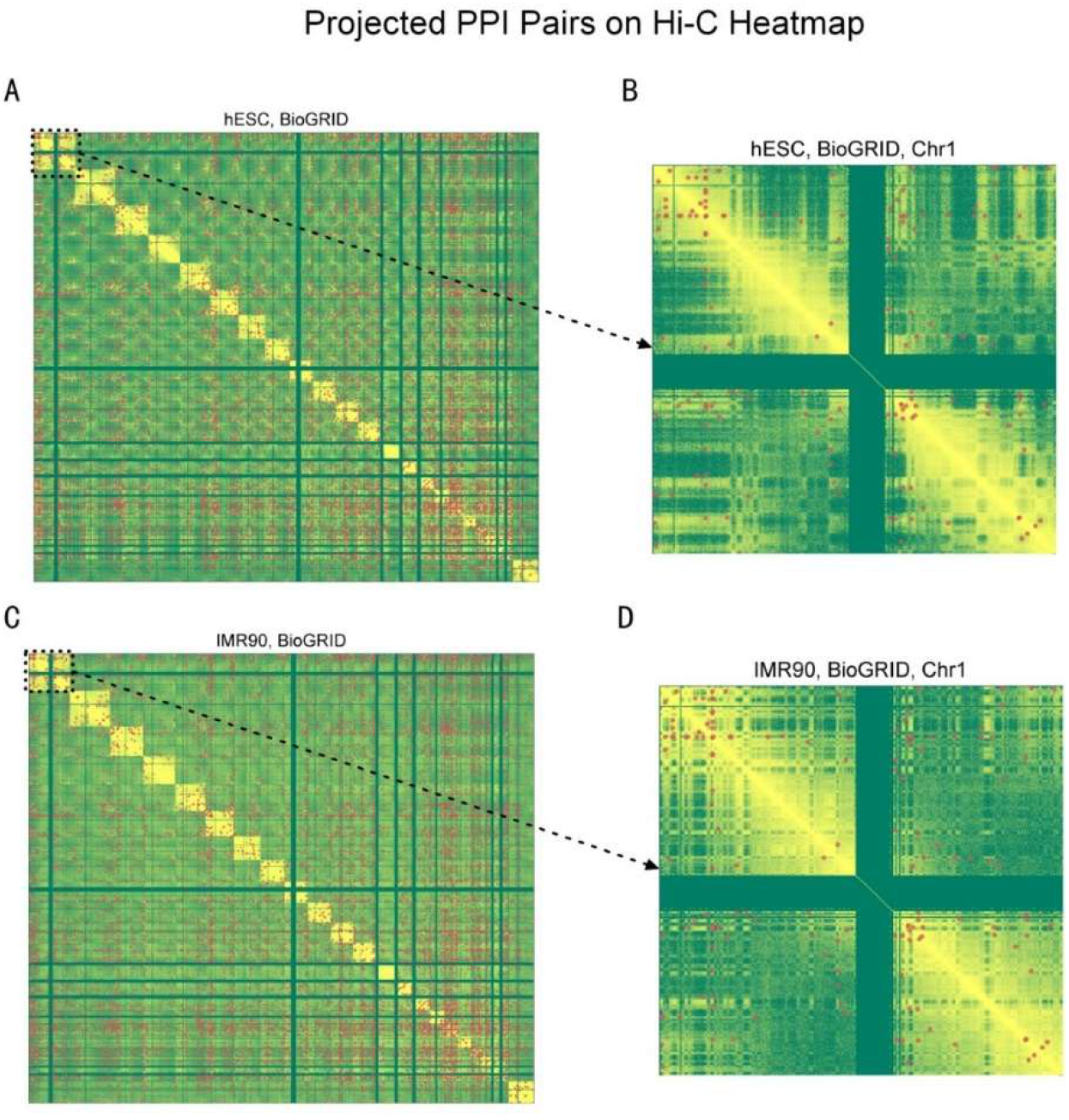
Gene-gene pair projections of PPIs overlaid on Hi-C heatmaps. A, B: PPIs from BioGRID database overlaid on whole genome and chromosome 1 of hESC Hi-C heatmaps respectively. C, D: PPIs from BioGRID database overlaid on whole genome and chromosome 1 of IMR90 Hi-C heatmaps respectively.

### Role of signal peptide in 3D genome assisted PPI

We believe that the above discovery can be explained by a logical conjecture that when two closely related genes that were once linearly next to each other in evolutional-wise lower complex genome, tend to be linearly separated far away or even re-located at different chromosome during evolution, to avoid linear-space batch error scenario such as replication or transcription. Yet, to still be able to cooperate, they remain spatially proximate so that their co-expression lead to co-localization of their RNAs and proteins counterparts, which further lead to protein-protein interaction. Additional pieces of patches can further enrich this conjectural storytelling picture and the signal peptide is one of them, as many proteins re-localization are guided by signal peptides.

To examine whether signal peptide affects the relation between 3D genome and PPI, we first labelled all the PPIs with at least one protein whose re-localization is assisted by signal peptide, and then we partition PPIs into two categories, i.e., signal peptide assisted PPI (SigPep PPI) and no signal peptide assisted PPI (Non-SigPep PPI). As Figure 5 indicates, gene-gene contact frequencies of PPIs that are not assisted by signal peptides tend to be higher than the gene-gene contact frequencies of PPIs that are assisted by signal peptides. This can be explained that for the interacting proteins that are brought together by signal peptides, their gene counterparts can be more freely located on the 3D genome, with larger spatial distances.

**Figure 5.**
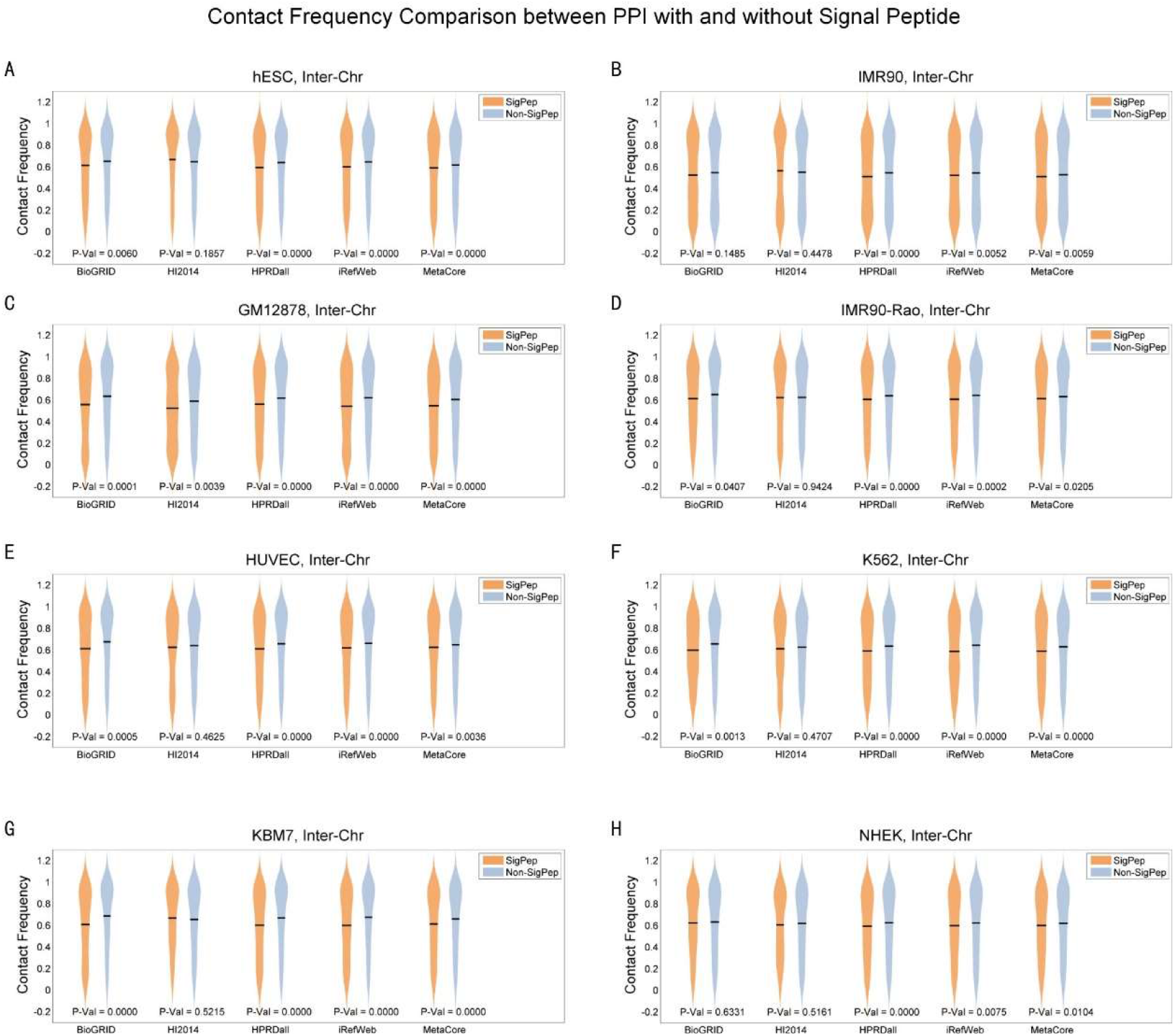
Comparison of contact frequencies between PPI with signal peptide assisted protein-protein co-localization and PPI without signal peptide assisted protein-protein co-localization.

### Applying 3D genome information in PPI prediction

Having the important discovery above, we then investigate whether adopting 3D genome information can contribute to more accurate PPI predictions, as none of the existing PPI prediction method ever consider PPI in the 3D genome perspective. We selected six representative PPI prediction methods and performed 5-fold cross validation on the PPI datasets, with and without 3D genome information. As Table 1 shows, the prediction accuracy in terms of AUC can be significantly improved if 3D genome information is employed.

**Table 1:** AUC performance of benchmark models without/with 3D positional information on all 4 datasets

| Dataset | AC-SVM* | CNN* | E-ELM* | AC-SAE* | DNN* | MCD-SVM* |
| --- | --- | --- | --- | --- | --- | --- |
| **BioGRID** | 0.7435/<br>**0.7586** | 0.7252/<br>**0.7822** | 0.7918/<br>**0.8129** | 0.8473/<br>**0.8694** | 0.8642/<br>**0.8962** | 0.9168/<br>**0.9174** |
| **HI2014** | 0.7589/<br>**0.7771** | 0.8690/<br>**0.8943** | 0.9192/<br>0.9228 | 0.8926/<br>**0.9156** | 0.9408/<br>**0.9449** | 0.9862/<br>**0.9865** |
| **iRefWeb** | 0.7310/<br>**0.7528** | 0.7499/<br>**0.8093** | 0.7623/<br>0.7862 | 0.8613/<br>**0.8785** | 0.8980/<br>**0.9017** | 0.9508/<br>**0.9518** |
| **MetaCore** | 0.7130/<br>**0.7380** | 0.7588/<br>**0.8140** | 0.7429/<br>0.7723 | 0.8614/<br>**0.8896** | 0.9114/<br>**0.9267** | 0.9537/<br>**0.9545** |

\*Performance measures the averaged AUC score on specific dataset in the order of: AUC without 3D genome information /AUC with 3D genome information

## Methods

### PPI and 3D genome data

We collected and curated five representative PPI datasets, namely BioGRID^19^, HI2014^20^, HPRDall^21^, iRefWeb^22^, and Clarivate MetaCore. The positive samples are interacting protein-protein pairs and the negative samples are draw from all the non-PPIs with different subcellular locations.

For the 3D genome data, we collected eight Hi-C datasets, namely hESC, IMR90, GM12878, HUVEC, IMR90-Rao, NHEK, K562, and KBM7^23,24^. The datasets are normalized using the KRNorm method and are curated so that intra-chromosomal heatmaps are of 40kb bin resolution and the inter-chromosomal heatmaps are of 500kb bin resolution.

### Encoding scheme and prediction methods

We selected six representative machine learning PPI prediction methods that are developed recently; each method has its own protein-protein encoding scheme. These methods are Auto-Covariance SVM (AC-SVM)^25^, Convolutional Neural Network (CNN)^26^, Ensemble Extreme Learning Machine (E-ELM)^27^, Auto-Covariance Stacked Encoder (AC-SAE)^28^, Deep Neural Network (DNN)^29^, and Multi-scale Continuous and Discontinuous SVM (MCD-SVM)^30^. We re-implemented all the six methods with different encoding schemes described in the references. To add 3D genome information, we computationally modeled 3D genome based on the Hi-C heatmaps and compute for each bin (500kb) a <x, y, z> coordinate each protein in the PPI datasets are assigned to a bin so that each protein has <x, y, z> feature 3-tuple.

## Discussion

In this work, we retrospect the DNA origins of the interacting proteins in the context of 3D genome and discovered that 1) if a gene pair is more proximate in 3D genome, their corresponding proteins are more likely to interact. 2) signal peptide involvement of PPI affects the corresponding gene-gene proximity in 3D genome space. 3) by incorporating 3D genome information, existing PPI prediction methods can be further improved in terms of accuracy. Combining our previous discoveries, we conjecture the existence of cellular compartmentalization driven by the chromatin 3D conformation. The concept of 3D genome driven cellular compartmentalization can well explain the co-localization of DNA elements lead to increased probability of the co-localization of their downstream elements including RNAs, proteins, and even metabolic molecules. More detailed investigation is needed to either further prove the 3D genome driven compartmentalization theory or utilize this theory in assisting 3D genome related researches.

## References

1 Zahiri, J., Bozorgmehr, J. H. & Masoudi-Nejad, A. Computational Prediction of Protein-Protein Interaction Networks: Algo-rithms and Resources. Curr Genomics 14, 397–414, doi:10.2174/1389202911314060004 (2013).

2 Venkatesan, K. et al. An empirical framework for binary interactome mapping. Nat Methods 6, 83–90, doi:10.1038/nmeth.1280 (2009).

3 Bonetta, L. Protein-protein interactions: Interactome under construction. Nature 468, 851–854, doi:10.1038/468851a (2010).

4 Ito, T. et al. A comprehensive two-hybrid analysis to explore the yeast protein interactome. Proc Natl Acad Sci U S A 98, 4569–4574, doi:10.1073/pnas.061034498 (2001).

5 Gavin, A. C. et al. Functional organization of the yeast proteome by systematic analysis of protein complexes. Nature 415, 141–147, doi:10.1038/415141a (2002).

6 van den Berg, D. L. et al. An Oct4-centered protein interaction network in embryonic stem cells. Cell Stem Cell 6, 369–381, doi:10.1016/j.stem.2010.02.014 (2010).

7 Shoemaker, B. A. & Panchenko, A. R. Deciphering protein-protein interactions. Part II. Computational methods to predict protein and domain interaction partners. PLoS Comput Biol 3, e43, doi:10.1371/journal.pcbi.0030043 (2007).

8 Tuncbag, N., Kar, G., Keskin, O., Gursoy, A. & Nussinov, R. A survey of available tools and web servers for analysis of protein-protein interactions and interfaces. Brief Bioinform 10, 217–232, doi:10.1093/bib/bbp001 (2009).

9 Li, X., Wu, M., Kwoh, C. K. & Ng, S. K. Computational approaches for detecting protein complexes from protein interaction networks: a survey. BMC Genomics 11 Suppl 1, S3, doi:10.1186/1471-2164-11-S1-S3 (2010).

10 Skrabanek, L., Saini, H. K., Bader, G. D. & Enright, A. J. Computational prediction of protein-protein interactions. Mol Biotechnol 38, 1–17, doi:10.1007/s12033-007-0069-2 (2008).

11 Raman, K. Construction and analysis of protein-protein interaction networks. Autom Exp 2, 2, doi:10.1186/1759-4499-2-2 (2010).

12 Santoni, D., Castiglione, F. & Paci, P. Identifying Correlations between Chromosomal Proximity of Genes and Distance of Their Products in Protein-Protein Interaction Networks of Yeast. Plos One 8, doi:ARTN e5770710.1371/journal.pone.0057707 (2013).

13 Lieberman-Aiden, E. et al. Comprehensive mapping of long-range interactions reveals folding principles of the human genome. Science 326, 289–293, doi:10.1126/science.1181369 (2009).

14 Rao, S. S. P. et al. A 3D Map of the Human Genome at Kilobase Resolution Reveals Principles of Chromatin Looping (vol 159, pg 1665, 2014). Cell 162, 687–688, doi:10.1016/j.cell.2015.07.024 (2015).

15 Fullwood, M. J. & Ruan, Y. ChIP-based methods for the identification of long-range chromatin interactions. J Cell Biochem 107, 30–39, doi:10.1002/jcb.22116 (2009).

16 Shi, Y. et al. Chromatin accessibility contributes to simultaneous mutations of cancer genes. Sci Rep 6, 35270, doi:10.1038/srep35270 (2016).

17 Akdemir, K. C. et al. Somatic mutation distributions in cancer genomes vary with three-dimensional chromatin structure. Nat Genet 52, 1178–1188, doi:10.1038/s41588-020-0708-0 (2020).

18 Shi, Y. et al. DeepAntigen: A Novel Method for Neoantigen Prioritization via 3D Genome and Deep Sparse Learning. Bioinformatics, doi:10.1093/bioinformatics/btaa596 (2020).

19 Oughtred, R. et al. The BioGRID interaction database: 2019 update. Nucleic Acids Res 47, D529–D541, doi:10.1093/nar/gky1079 (2019).

20 Ideker, T. & Valencia, A. Bioinformatics in the human interactome project. Bioinformatics 22, 2973–2974, doi:10.1093/bioinformatics/btl579 (2006).

21 Keshava Prasad, T. S. et al. Human Protein Reference Database--2009 update. Nucleic Acids Res 37, D767–772, doi:10.1093/nar/gkn892 (2009).

22 Turner, B. et al. iRefWeb: interactive analysis of consolidated protein interaction data and their supporting evidence. Database (Oxford) 2010, baq023, doi:10.1093/database/baq023 (2010).

23 Dixon, J. R. et al. Topological domains in mammalian genomes identified by analysis of chromatin interactions. Nature 485, 376–380, doi:10.1038/nature11082 (2012).

24 Rao, S. S. et al. A 3D map of the human genome at kilobase resolution reveals principles of chromatin looping. Cell 159, 1665–1680, doi:10.1016/j.cell.2014.11.021 (2014).

25 Guo, Y., Yu, L., Wen, Z. & Li, M. Using support vector machine combined with auto covariance to predict protein-protein interactions from protein sequences. Nucleic Acids Res 36, 3025–3030, doi:10.1093/nar/gkn159 (2008).

26 Lecun, Y., Bottou, L., Bengio, Y. & Haffner, P. Gradient-based learning applied to document recognition. P Ieee 86, 2278–2324, doi:Doi 10.1109/5.726791 (1998).

27 You, Z. H., Lei, Y K., Zhu, L., Xia, J. F. & Wang, B. Prediction of protein-protein interactions from amino acid sequences with ensemble extreme learning machines and principal component analysis. Bmc Bioinformatics 14, doi:Artn S1010.1186/1471-2105-14-S8-S10 (2013).

28 Sun, T. L., Zhou, B., Lai, L. H. & Pei, J. F. Sequence-based prediction of protein protein interaction using a deep-learning algorithm. Bmc Bioinformatics 18, doi:ARTN 27710.1186/s12859-017-1700-2 (2017).

29 Wainberg, M., Merico, D., Delong, A. & Frey, B. J. Deep learning in biomedicine. Nat Biotechnol 36, 829–838, doi:10.1038/nbt.4233 (2018).

30 You, Z. H. et al. Prediction of protein-protein interactions from amino acid sequences using a novel multi-scale continuous and discontinuous feature set. Bmc Bioinformatics 15, doi:ArtnS910.1186/1471-2105-15-S15-S9 (2014).

